## Supplemental Material for "S1 hydrophobic residues modulate voltage sensing phosphatase enzymatic function and voltage sensing"

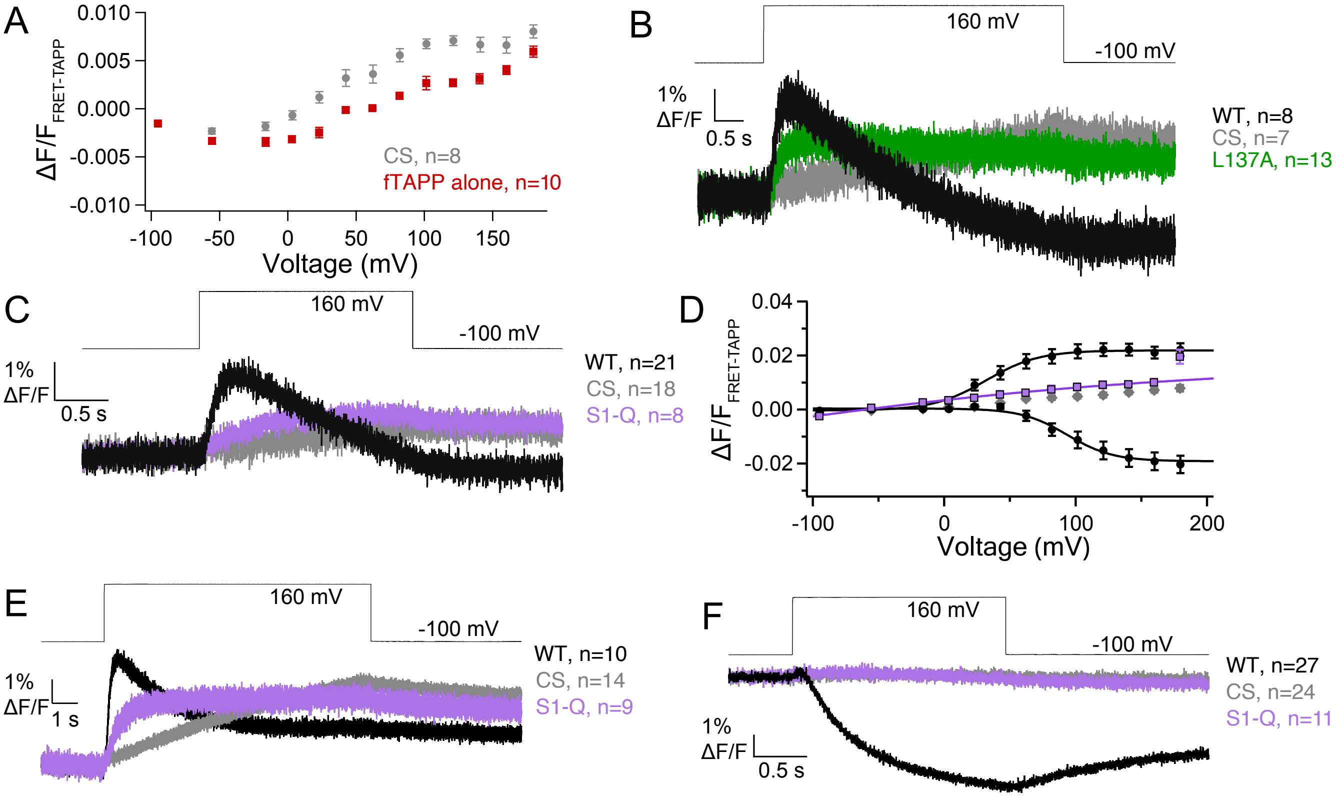


Supplemental Figure 1: A) Voltage dependent fTAPP activity in *X. laevis* oocytes. Cells co-expressing catalytically inactive Ci-VSP (C363S, CS) with fTAPP or fTAPP alone. Significantly more background activity from *X. laevis* VSP2 (Xl-VSP2) is observed in cells expressing inactive Ci-VSP. This result suggested Xl-VSP2 may be more efficiently trafficked to the plasma membrane in the presence of Ci-VSP. B) Averaged data for a long voltage step (5s) from -100 mV to 160 mV for L137A, WT and CS with the fTAPP biosensor. Unsubtracted data from Fig. 3D. While the CS protein is inactive, the resulting fTAPP increase is significant, indicating a substantial amount of Xl-VSP2 activity for the PI(3,4,5)$P_3$ to PI(3,4)$P_2$ reaction. C) Averaged data for a voltage step from -100 mV to 160 mV (2s) for S1-Q. 5-phosphatase activity is dramatically reduced while 3-phosphatase activity appears eliminated. D) Voltage dependent fTAPP activity for S1-Q, WT and CS from the 2s data. While S1-Q is still active, the activity is almost linear and barely above the CS control. E) Averaged data for a long voltage step (10s) from -100 mV to 160 mV for S1-Q, WT and CS with the fTAPP biosensor. The CS protein shows a significant degree of Xl-VSP activity at the longer time scale. Unsubtracted data from Fig. 3F. F) Averaged data for a 2s voltage step from -100 mV to 160 mV for the S1-Q mutation with the fPLC biosensor. S1-Q activity above background (CS) was not observed.

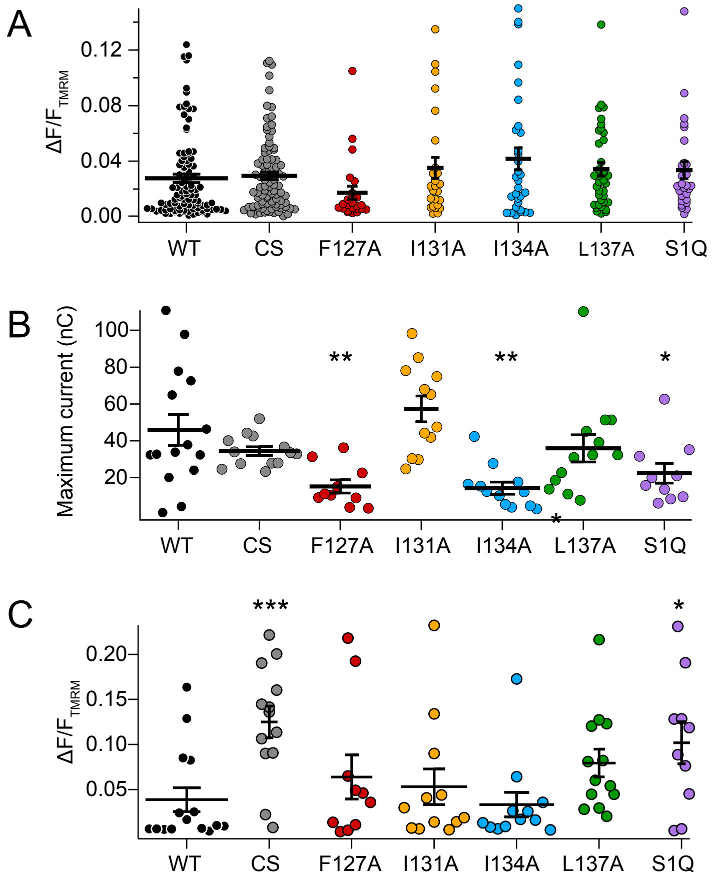


Supplemental Figure 2: A) VCF expression data for all the cells used in the activity assays. No statistically significant difference was found between WT and the mutations. WT n= 97, CS n=95, F127A n=24, I131A n=25, I134A n=31, L137A n=38, S1Q n=27. B) Maximum off-sensing charge from a 150mV step. Lower current was found for F127A, I134A and S1-Q. WT n=15, CS n=13, F127A n=10, I131A n=12, I134A n=12, L137A n=13, S1Q n=10. C) VCF data for the same cells in B. Larger fluorescence was found for CS and S1-Q. Statistics of mutations vs WT determined using the Welch's t-test with a two-tailed distribution. *** $p < 0.001$; ** $p < 0.01$; * $p < 0.05$

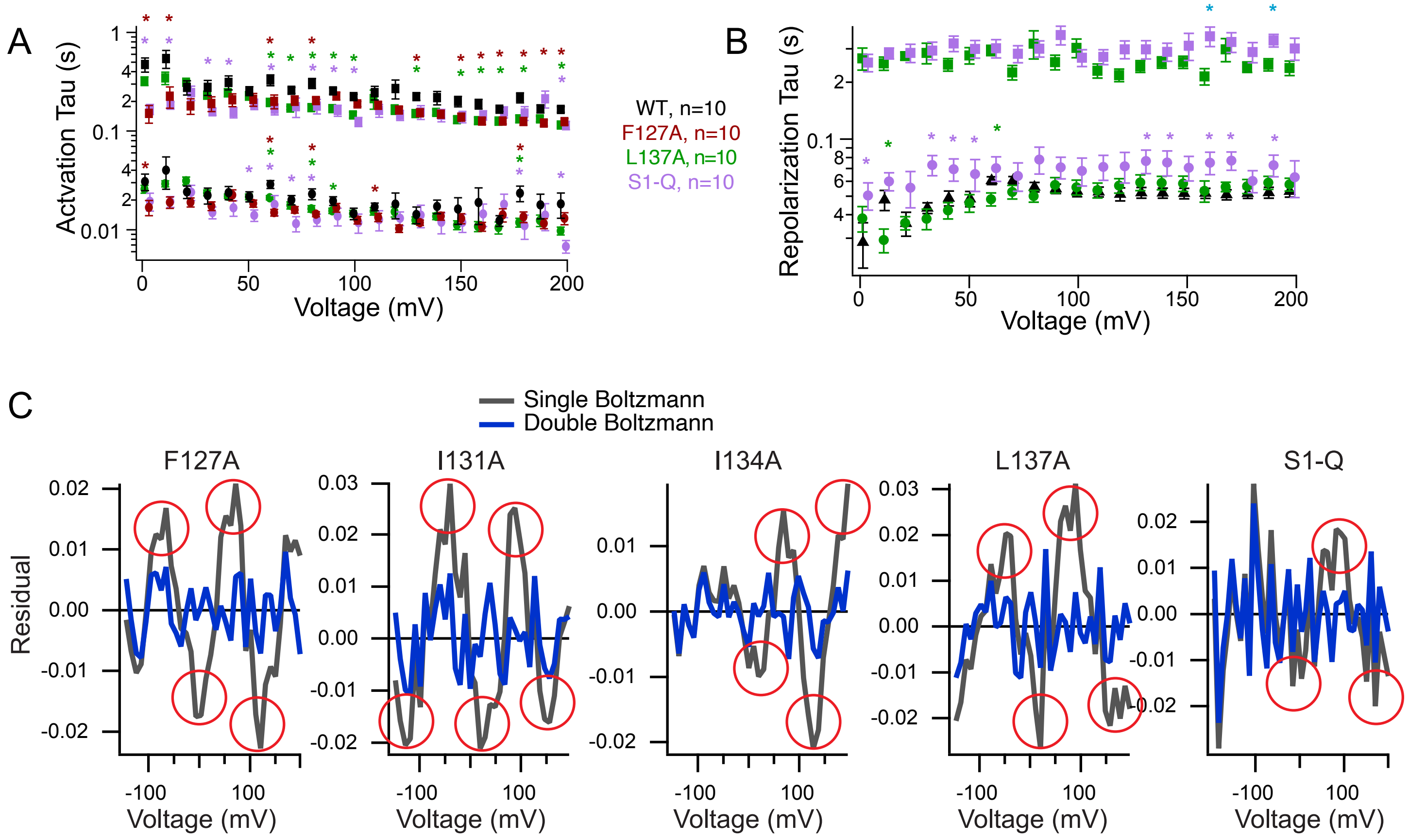


Supplemental Figure 3: A) Activation kinetic analysis for WT, F127A, L137A and S1-Q VCF. Data from Fig 6A and 6C were fit with double exponential fits and the resulting $\tau_{a1}$ and $\tau_{a2}$ values plotted versus the voltage measured for each trace. Student t-tests *p=0.001-0.049 shown against WT in the corresponding color for each mutation. B) Repolarization kinetics were fit with a single exponential (WT) or a double exponential (L137A, S1-Q). The resulting $\tau_{r1}$ and $\tau_{r2}$ values were plotted versus the corresponding voltage. Student t-tests *p=0.0025-0.034 shown against WT in the corresponding color for each mutation, except for $\tau_{r2}$ values which are shown in cyanand are L137A versus S1-Q. C) Residual for each of the individual mutations and S1-Q, comparing the single and double fits. Significant discrepencies in the single Boltzmann sigmoids are circled in red. The double Boltzmann sigmoid consistently fits all the mutant VCF data better (fits from Fig 5B, D).

Supplementary Table 1: Primers

| | |
|---|---|
| **His forward** | GGGGATCCGCCACCATGCATCATCACCACCACCATGAGGGATTCGACGGT TC |
| **His reverse** | GAACCGTCGAATCCCTCATGGTGGTGGTGATGATGCATGGTGGCGGATCCCC |
| **FLAG forward** | GGGGATCCGCCACCATGGACTACAAAGACGATGACGACAAGGAGGGATTCGACGGTTC |
| **FLAG reverse** | GAACCGTCGAATCCCTCCTTGTCGTCATCGTCTTTGTAGTCCATGGTGGCGGATCCCC |
| **G214C forward** | GAAACAGGAGCCGATTGTTTGGGGAGATTG |
| **G214C reverse** | CAATCTCCCCAAACAATCGGCTCCTGTTTC |
| **C363S forward** | GATAGCGATTCACTCTAAAGGCGGGAAG |
| **C363S reverse** | CTTCCCGCCTTTAGAGTGAATCGCTATC |
| **F127A forward** | GGAGTCTTCCTAATTGCATTGGACATCATCCTCATG |
| **F127A reverse** | CATGAGGATGATGTCCAATGCAATTAGGAAGACTCC |
| **I131A forward** | CTAATTTTCTTGGACATCGCACTCATGATCATTGATC |
| **I131A reverse** | GATCAATGATCATGAGTGCGATGTCCAAGAAAATTAG |
| **I134A forward** | CATCATCCTCATGGCAATTGATCTCAGTC |
| **I134A reverse** | GACTGAGATCAATTGCCATGAGGATGATG |
| **L137A forward** | CTCATGATCATTGATGCCAGTCTTCCAGGAAAAAGTG |
| **L137A reverse** | CACTTTTTCCTGGAAGACTGGCATCAATGATCATGAG |
| **F127A I131A forward** | GTCTTCCTAATTGCATTGGACATCGCACTCATGATC |
| **F127A I131A reverse** | GATCATGAGTGCGATGTCCAATGCAATTAGGAAGAC |
| **I134A L137A forward** | CTCATGGCAATTGATGCCAGTCTTCCAGGAAAAAGTG |
| **I134A L137A reverse** | CACTTTTTCCTGGAAGACTGGCATCAATTGCCATGAG |

Supplementary Table 1: VCF, kinetics of S4 motions from 200 mV step

| | | Activation | | Repolarization | |
|---|---|---|---|---|---|
| | **n** | $\tau_{a1}$ | $\tau_{a2}$ | $\tau_{r1}$ | $\tau_{r2}$ |
| **WT** | 10 | 0.018 ± 0.005 | 0.17 ± 0.01 | 0.053 ± 0.004 | N/A |
| **F127A** | 10 | 0.013 ± 0.002 | 0.12 ± 0.01 * | 0.0602 ± 0.006 | N/A |
| **I131A** | 13 | 0.015 ± 0.003 | 0.15 ± 0.02 | 0.069 ± 0.004 | N/A |
| **I134A** | 15 | 0.010 ± 0.001 | 0.15 ± 0.01 | 0.066 ± 0.004 | N/A |
| **L137A** | 10 | 0.010 ± 0.001 | 0.12 ± 0.01 * | 0.058 ± 0.004 | 0.24 ± 0.03 |
| **S1Q** | 10 | 0.007 ± 0.001 * | 0.11 ± 0.02 * | 0.06 ± 0.01 | 0.30 ± 0.05 |

* Student's t-test $p < 0.05$
